# A remarkable degree of conformance between the visual streak of the Mongolian gerbil and the human central retina

**DOI:** 10.1101/2024.08.28.610039

**Authors:** Alexander Günter, Mohamed Ali Jarboui, Regine Mühlfriedel, Mathias W. Seeliger

**Affiliations:** Division of Ocular Neurodegeneration, Institute for Ophthalmic Research, Centre for Ophthalmology, University of Tübingen, Tübingen, Germany; Core Facility for Medical Bioanalytics (CFMB), University of Tübingen, Tübingen, Germany

**Keywords:** Mongolian gerbil, macula, visual streak, parafovea, age-related macular degeneration, retinal pigment epithelium, photoreceptors, phagocytosis, diurnal rodents

## Abstract

The Mongolian gerbil (MG) is a day-active rodent that lives in desert-like environments and thus very much relies on vision. To allow for a particularly exact view of the horizon, the region of highest visual acuity, the visual streak (VS), forms a horizontal band across the retina between the projection areas of the sky and the ground. Here, we assessed the retinal basis of this specialized region and compared the findings to the human central high-acuity region culminating in the macula. We found an increased density of cones, elongated photoreceptor outer segments (OSs) and a minimization of intersecting surface vessels to improve quality of vision, all of which is analogous to the human macula. Stunningly, also the area of retinal pigment epithelium (RPE) cells was significantly smaller in the VS region than in the periphery, again similar to what is found in the human macula. Our data therefore suggest that the remarkable degree of conformance between the VS of the MG and the human macula renders the MG a promising rodent, non-primate model of the central human retina.

## 1 Introduction

Human reading and color vision very much depends on the function of the cone photoreceptor system in the central retina at the back of the eye, in particular the macula. There is a wealth of inherited and acquired diseases that lead to a general or localized dysfunction of these sites, impairing regular vision up to legal blindness. Most commonly known is age-related macular degeneration (AMD), the leading cause of irreversible blindness in individuals over 50 years of age, a chronic and progressive disease characterized by the degeneration of photoreceptors and retinal pigment epithelium (RPE) cells in the macula. In AMD, a slow degeneration of the RPE cell layer has the accumulation of intracellular and extracellular deposits knows as lipofuscin and drusen, respectively, as hallmarks of the disease (Wong et al., 2022). The decrease in macular phagocytic activity of photoreceptor OSs by the RPE is directly related to the accumulation of these deposits in ageing and AMD (Inana et al., 2018). In order to investigate retinal physiology and pathophysiology and to develop therapeutic strategies, models are needed that have a high conformance with the human situation. Unfortunately, mice are currently the most common models in terms of genetic manipulation to produce homologous disease genotypes, but are –due to their nocturnal lifestyle-physiologically not optimally suited to study diseases of the cone system. In this work, we have assessed whether the Mongolian gerbil (MG), a diurnal rodent originating from an environment of semi-deserts and steppes (Scheibler & Waiblinger, 2018), may turn out to be superior to mice as a model for human central retina.

The mammalian retina uses rod and cone photoreceptors as primary sensory cells for the detection of light. Rods and cones translate light stimuli into electrical signals, which travel along the retinal network to the brain (Kolb, 2003). Rods are most sensible to light stimuli in dim light (scotopic) conditions, while cones provide high-acuity and color vision in brighter (photopic) conditions. Photoreceptors are supported by the RPE, a monolayer of cells that forms the blood-retinal barrier (Lakkaraju et al., 2020). The RPE provides important support to photoreceptors, including regulation of vitamin A supply, absorbance of excess light by melanin, and participation in the daily renewal of ROS. RPE dysfunction may thus lead to retinal degeneration (Kevany & Palczewski, 2010).

The human macula is a specialized region for high-acuity vision with a diameter of 3-5.5 mm (Hussey et al., 2022). At the center of the macula, the fovea is located, featuring densely packed cones with elongated outer segments (OSs), but no rods. Adjacent to the fovea follow the para- and perifovea, ring-shaped cone-rich regions that form the ‘macular shoulder’ characterized by a continuous increase in rod and a decrease in cone cell density with eccentricity. Below the macula, the area of supporting RPE cells is smaller and they are also taller when compared to more peripheral regions (Ortolan et al., 2022; Weiter et al., 1986). Above the macula, there is also a lack of retinal vessels intersecting the path of light to avoid any interference with vision (Ahnelt & Kolb, 2000; Hussey et al., 2022; Kolb, 2005; Seeliger et al., 2005).

Retinal organization of species often arises from their habitat and their role in the animal community (e.g. predator vs. prey). The evolutionary pressure to effectively collect the visual information that fits the specific need for survival and reproduction may lead to major differences in retinal topography (Baden et al., 2020; Schiviz et al., 2008). Mice live in a nocturnal environment, and therefore lack any region of high-acuity vision throughout the retina. Humans on the other hand are diurnal and rely on binocular vision that stimulated the development of a very advanced cone system organization (i.e., macula and fovea). In many mammalian species, a horizontal VS is found, which is ideal for habitats with visual scenes dominated by the horizon and a clear separation of sky and ground. Predators with a higher demand for acuity often have developed an additional *area centralis* within or adjacent to the VS (Hauzman et al., 2018; Rapaport & Stone, 1984). Examples for these different retinal organization types are given in Fig. 1.

**Figure 1.**
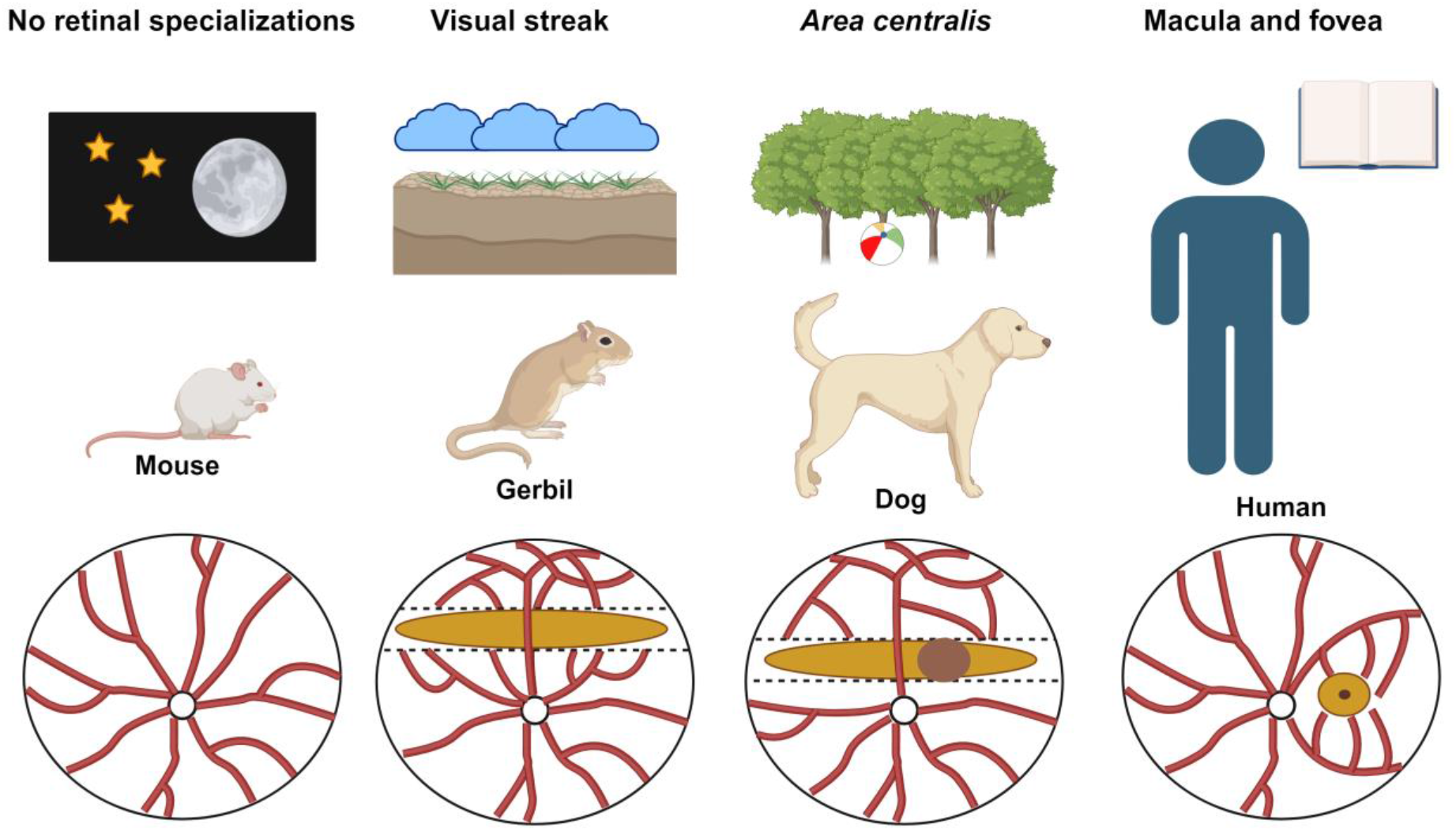
Examples of different retinal organizations. Mice are nocturnal animals and therefore require a high degree of light sensitivity (provided by rods) but no site of high-acuity vision (provided by cones). The absence of any high-acuity region may be observed as a rather homogeneous build-up of the murine retina (left). Diurnal rodents like the Mongolian gerbil (MG) originate from the semi-deserts and steppes of central asia and use a visual streak (VS) to scan the horizon for potential predators and their own flock. To enhance image quality, intersecting blood vessels that would optically interfere are reduced to a minimum (second to left). Some predators like dogs and cats have an additional ring-shaped area of high visual acuity within or close by the VS, the *area centralis*. These areas allow for some binocular vision that cannot be obtained by superposition of VSs. Humans that heavily rely on binocular vision have a very high-resolution area, the macula, with a central rod-free fovea. White circle: optic nerve head; red lines: major retinal vessels; dark yellow bar: VS; dark brown circle: area centralis; dark yellow circle with black dot: macula with central fovea.

Mice are currently the most widely used animal models for research on retinal diseases due to the presence of well-established genetically matched disease models. While major advances in the understanding of rod-based disease mechanisms have been feasible, research on cone-based disorders has been limited by the anatomically and physiologically substantially aberrant cone system when compared to humans. Although the fraction of cones within all photoreceptors (about 3-5%) is similar to the human ratio, their build-up and topographical organization is very different as shown above. As nocturnal species, high acuity vision is of low importance, and so there is no horizontal band like a VS or even a macula. Most importantly, the majority of cones co-express the cone-specific optical pigments S- and M-opsin. These opsins follow counter-directional dorsoventral gradients, i.e. M-opsin is found in highest levels in the dorsal region and expression declines towards the ventral end of the retina, while S-opsin is highly expressed in cones of the ventral retina and its levels decline towards the dorsal end (Applebury et al., 2000; Carter-Dawson & LaVail, 1979; Wang et al., 2011). In contrast, there is only one type of opsin per cone expressed in both human and MG retina and the expression of the different opsins does not follow a similar gradient (Günter et al., 2024). Although the VS of the MG is not a rod-free region like the very central fovea, it still shares some important characteristics similar to the parafoveal region. The parafovea has been identified as the region where initial pathological changes are found in a number of disorders including AMD, and so an animal model of this area would be of great relevance (Curcio et al., 1996; Sarks et al., 1988). For these reasons, it was investigated in this study whether the MG may satisfy the high unmet need for small animal models with a cone system organization better matching the human situation.

## 2 Results

### The visual streak of the Mongolian gerbil: Cone density and rod-to-cone ratio

A key feature of high-acuity vision is an increased density of cone photoreceptors in respective topographical areas. In contrast, rod photoreceptor density does usually not vary to a similar degree as spatial resolution is less relevant than sensitivity in dim light conditions. Accordingly, a substantially increased density of cones is present in human foveal and parafoveal regions when compared to the periphery (Curcio et al., 1990). Our data show that this is also true for the VS of the MG. We found a cone density of 50660 ± 3370 cones / mm^2^ in the VS, much higher than the densities of 37743 ± 2054 cones / mm^2^ and 28476 ± 1696 cones / mm^2^ in the near and far periphery, respectively (Fig. 2A, B). In contrast, we recorded a rod density of 345014 ± 15774 rods / mm^2^ in the VS, 339391 ± 11337 rods / mm^2^ in the near periphery, and 312946 ± 10978 rods / mm^2^ in the far periphery (Fig. 2C; Table 1). These distribution characteristics are in excellent agreement with a high-acuity topography of the retina and very much dislike that found in mice.

**Table 1.** Comparison of photoreceptor and retinal pigment epithelium (RPE) characteristics of Mongolian gerbils and humans.

| Mongolian gerbil (MG) | Rod count per mm<sup>2</sup> (x1000) | Cone count per mm<sup>2</sup> (x1000) | Rod-to-cone ratio (RCR) | RPE cell area in μm<sup>2</sup> |
| --- | --- | --- | --- | --- |
| Visual streak (VS)<sup>(1)</sup> | **354**<br>[328-357] | **51**<br>[47-54] | **6.84**<br>[6.64-6.98] | **317.9**<br>[294.9-329.1] |
| Far periphery<sup>(1)</sup> | **313**<br>[304-322] | **29**<br>[27-30] | **11**<br>[10.55-11.47] | **442.1**<br>[392.5-467.2] |
| Human | Rod count per mm<sup>2</sup> (x1000)<sup>(2)</sup> | Cone count per mm<sup>2</sup> (x1000)<sup>(2)</sup> | Rod-to-cone ratio (RCR)<sup>(2)</sup> | RPE cell area in μm<sup>2</sup><sup>(3,4)</sup> |
| Parafovea (1.5 mm eccentricity) | **100** | **15** | **7** | **159.705**<br>± 18.46 |
| Far periphery (20 mm eccentricity) | **50** | **5** | **10** | **331.874**<br>± 27.23 |

<sup>(1)</sup> **Median [25-75% quantiles]** <sup>(2)</sup> **Human photoreceptor data from (Curcio et al., 2024; Curcio et al., 1990)** <sup>(3)</sup> **Human RPE data from (Ortolan et al., 2022)** <sup>(4)</sup> **Median ± SD**

**Figure 2.**
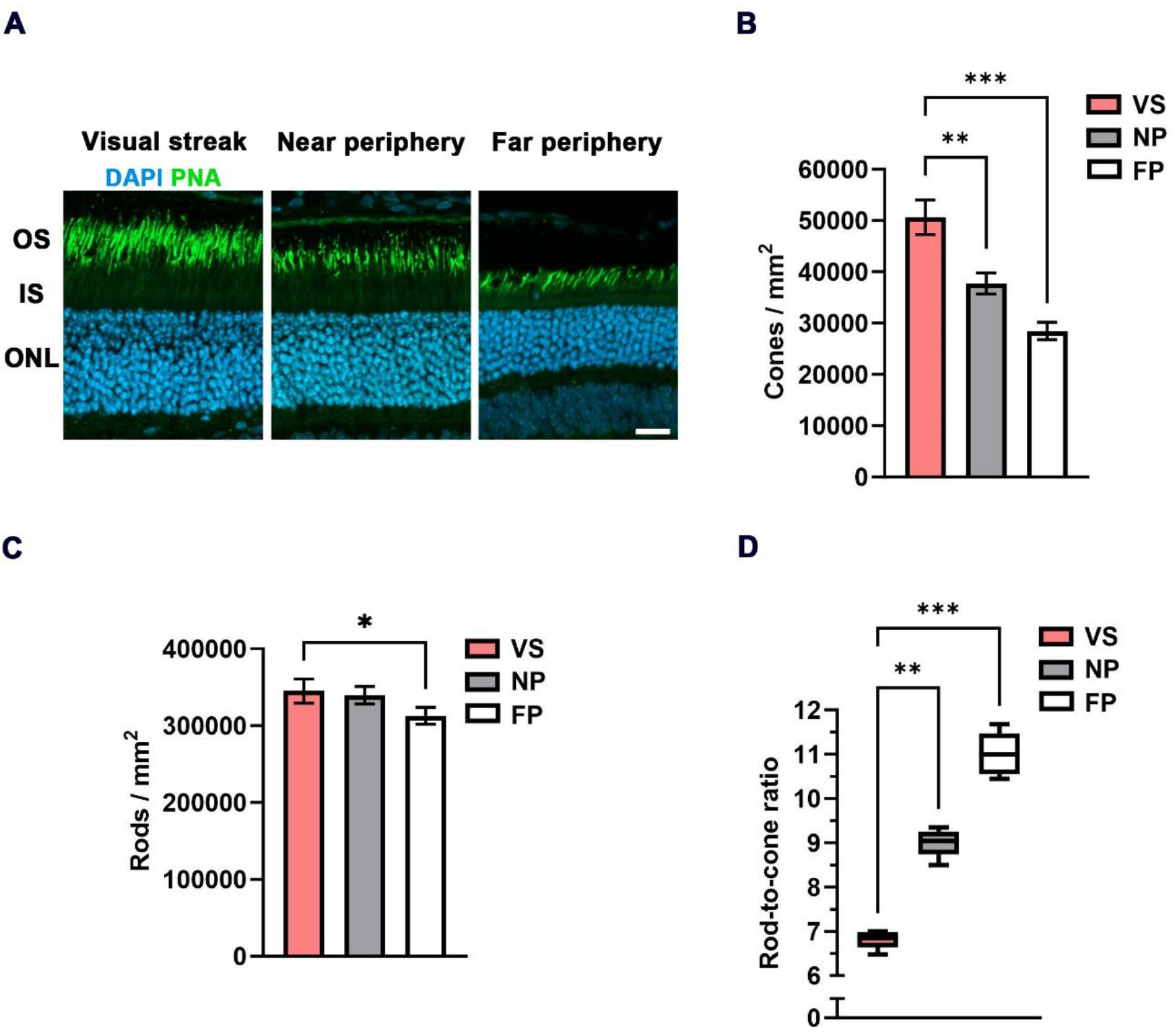
Densities of photoreceptors and RCR in the VS of the MG. **(A)** Cone outer segment (OS) immunohistochemistry (PNA-based) indicates an increase in OS length in the VS when compared to the near and far peripheral regions. DAPI was used as nuclear counterstaining and for counting of photoreceptors in the outer nuclear layer (ONL). IS = Inner segment; OS = Outer segment; Scale bar = 20 μm. **(B)** Quantification of the density of cones / mm^2^ in the VS compared to the near and far peripheral retinas (n = 5 eyes). **(C)** Quantification of the density of rods / mm^2^ µm in the VS compared to the near and far peripheral retinas (n = 5 eyes). Error bars represent the standard deviation. **(D)** Quantitative evaluation of the RCR (box-and-whisker-plot) in the VS compared to the near and far peripheral retinas (n = 5 eyes). Boxes: 25%-75% quantile range, whiskers: 5% and 95% quantiles, central line: median; * = p<0.05, ** = p<0.01 and *** = p<0.001. VS = Visual streak, NP = Near periphery, FP = Far periphery.

A further common finding in the human foveal region and the parafovea is the elongation of the cone OSs (Wang et al., 2023). In the VS of the MG, we also observed a comparable elongation of cone OSs when compared to regions in the peripheral retina (Fig 2A). This likewise applies to rod cells (Günter et al., 2024) to allow for physical contact to the supporting RPE. A proteomic differential expression analysis between the VS and the peripheral retina/RPE interface was performed to obtain further insight in possible changes in protein expression. In this analysis, we found an enhanced abundance of rod OS markers like rhodopsin and cone OS markers like M-opsin in the VS (Supplementary Figure 2A). A list of all proteins identified and quantified across VS and periphery are presented in the Supplementary Spreadsheet 1.

An important marker amalgamating the distribution topography of rods and cones is the rod-to-cone ratio (RCR). The fraction is formed that way as rod density may become zero (in the fovea), but cone density does not, and so rod density is not suitable as denominator. This means, the RCR is low in regions of high acuity, and high in less specialized areas. In the VS of the MG, we found an RCR of 6.8 ± 0.2, significantly lower than the ratios of 9 ± 0.31 in the near periphery and 11 ± 0.48 in the far periphery (Fig. 2D). A comparison to the human data given in table 1 shows a RCR ratio of 7 at an eccentricity of 1.5 mm (parafovea), which is very similar to the value observed in the VS of the MG. Further, the RCR of 10 in the far peripheral region of the human retina is also comparable to the corresponding region in the MG retina.

### The visual streak of the Mongolian gerbil: RPE cell area and shape

In addition to the conformance between the VS of the MG and the human central retina in terms of photoreceptor morphology and distribution, we moved on to an analysis of morphology and distribution of the underlying RPE cells. In human studies, it has been observed that the horizontal extensions (and thus the area) of RPE cells in the central retina are smaller than in the periphery (Dorey et al., 1989; Ortolan et al., 2022; Roorda et al., 2007) (Table 1). In the MG, we recorded the distances between RPE cell nuclei as a measure of horizontal extension and calculated the respective average area per cell (Figs. 3A, B) based on a formula as shown in the Methods section and the Supplementary material (Supplementary Figure 1B).

**Figure 3.**
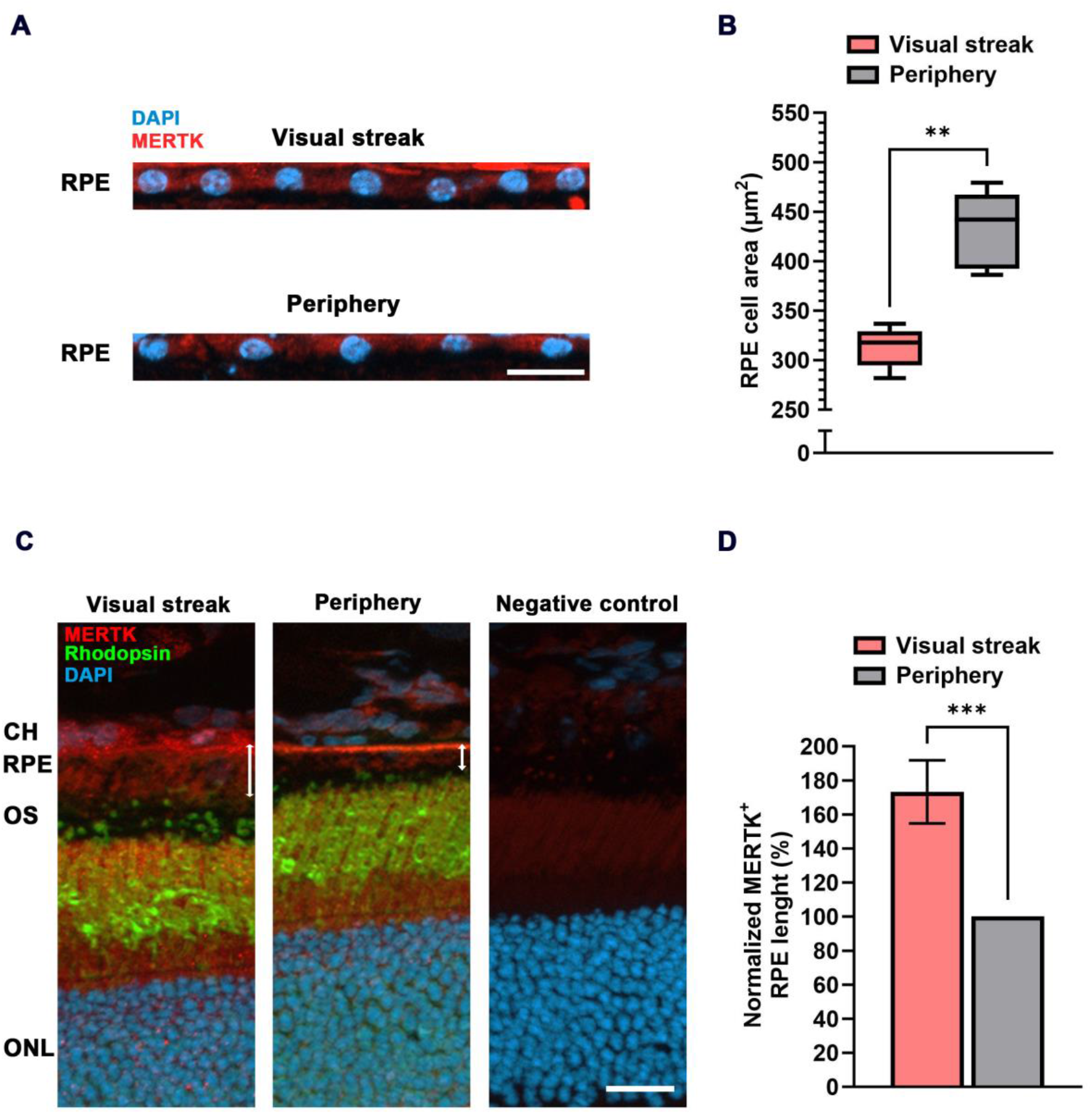
RPE cell characteristics in the VS of the MG. **(A)** Immunohistochemical staining of nuclei (DAPI) in the RPE monolayer of the VS (top) and the peripheral retina (bottom). MERTK was used to mark the RPE layer. RPE = Retinal pigment epithelium; Scale bar = 20 μm. **(B)** Quantitative evaluation of the cell area in μm^2^ (box-and-whisker-plot) in the VS compared to the peripheral retina (samples from n = 5 eyes). Boxes: 25%-75% quantile range, whiskers: 5% and 95% quantiles, central line: median. **(C)** Increased height of RPE cells (marked by MERTK staining) in the VS when compared to those in the periphery (white double arrows). CH = Choroid; OS = Outer segment; ONL = Outer nuclear layer; Scale bar = 20 μm. **(D)** Quantification of RPE height in the VS compared to the peripheral retina (n = 5). The error bar represents the standard deviation; ** = p<0.01 and *** = p<0.001.

RPE cells form a monolayer that is able to replace gaps by division of neighboring cells, and so a certain number of RPE cells is known to be binucleate (i.e. having two nuclei). We therefore checked the abundance of such binucleate cells in our samples and removed these from calculations of cell area. The average horizontal extension of RPE cells in the VS of the MG (19.16 ± 0.62 µm) was smaller in comparison to the 22.59 ± 1.01 µm found in the periphery (Fig. 3A; Supplementary Figure 1A). The extension data were used to calculate the cell area, resulting in an RPE cell area of 317.9 ± 20.34 µm^2^ in the VS, significantly smaller than the area of 442.1 ± 38.97 µm^2^ in the peripheral retina (Fig. 3B).

In addition to the reduced area, human RPE cells in the macula are also taller when compared to those in the periphery (Weiter et al., 1986). We found the same effect in the VS of the MG based on a Mer tyrosine kinase (MERTK) staining, indicating an increase of 73.3 ± 18.5 % in length compared to the peripheral retina (Figs. 3C, D). This increase in length (∼70%) is more than what would be needed to compensate for the reduced area (∼40%) in terms of an increase in RPE cell volume, suggesting an increased content of intracellular phagosomes containing material from photoreceptor OS tips (Fig. 3C). This is supported by a proteomic differential expression analysis between the VS and the peripheral retina/RPE interface, which confirmed an enhanced abundance of MERTK (Supplementary Figure 2A) together with non-membrane-bound RPE proteins that directly interact with MERTK for the engulfment of OS phagosomes like Gas6 and focal adhesion kinase (FAK) (Supplementary Figure 2B) (Finnemann, 2003). In line with these findings, OS and RPE proteins related to the visual cycle were also more abundant in the VS of the MG when compared to the peripheral retina (Supplementary Figures 2A, B).

## 3 Discussion

In this study, we assessed the degree of conformance between the VS of the MG and the human central retina as part of the search for a rodent, non-primate model system for central human retinal disorders. Specifically, we characterized photoreceptor density, RCRs, RPE cell extension, height, and area, between the VS and the retinal periphery. Additionally, we obtained proteomic profiles of retina and RPE in the VS of the MG and compared those to data from the peripheral retina. We found that the RCR in the VS of the MG is similar to the one found in the human parafovea. Also, similar to the human parafovea, the RPE cell area in the VS was smaller and cells taller in comparison to the peripheral retina. The increase in length (about 70%) exceeded the reduction in area (about 40%), indicating an increased volume of central RPE cells. Proteomic data supported these findings via an enhanced abundance of key proteins for photoreceptor OS phagocytosis and RPE visual cycle in the VS of the MG.

In terms of research for macular degenerations like AMD, the parafovea is an important area for understanding disease progression. This area has been described as particularly vulnerable during ageing, and as the beginning area of AMD (Curcio et al., 1993; Sarks et al., 1988). Before the occurrence of characteristic visible disturbances in the RPE cells, it is actually the cell death of mainly rods in the parafovea that was shown to influence the early stages of this disease. Together with this, rod sensitivity in this region is also severely affected (Curcio et al., 2024; Curcio et al., 1996). The VS of the MG showed the same RCR as in the human parafovea and could therefore be an interesting model to understand the pathophysiology of AMD. Moreover, the retina of the MG showed a similar periphery to center topographic transition of photoreceptor densities and the resulting RCR when compared to the human retina (Fig. 4). In comparison, mice do not show throughout their retina a RCR as low as observed in the parafovea or even the near periphery (Curcio et al., 2024; Volland et al., 2015), due to their species lifestyle (Baden et al., 2020).

**Figure 4.**
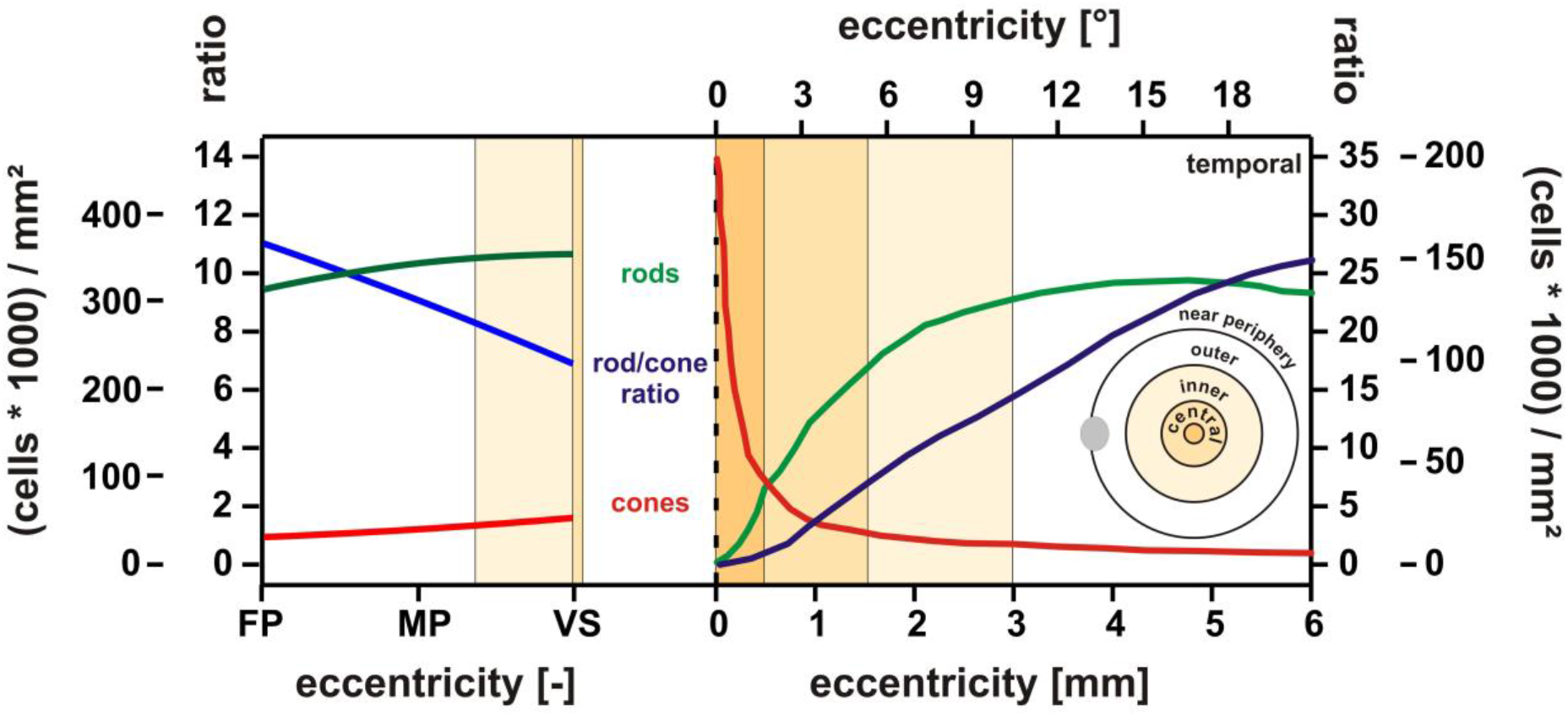
Degree of conformance of retinal photoreceptor densities and RCR between the MG and the human retina. Left: MG data for the three eccentricity regions VS, mid periphery (MP), and far periphery (FP). **Right:** Human eccentricity data redrawn from (Curcio et al., 2024), Figure 2. The courses of rod (green) and cone (red) photoreceptor density (scale (cells*1000)/mm^2^) and the resulting rod-to-cone ratio RCR (blue) indicate a similar topographic transition characteristic from the periphery (left end in the MG and right end in human) towards the retinal center, the VS in the MG (right end of left partial graph) and the human macula/fovea (left end of right partial graph). The VS data match the human situation at about 1.5-2 mm eccentricity, i.e. the parafoveal “macular shoulder” range.

To further understand the pathophysiology of macular diseases, a detailed assessment of the interaction of photoreceptors with RPE cells is of great importance. However, this appears to be dependent on a valid representation of macular RPE cells in a model system, as these are obviously different from peripheral ones. In order to address this aspect, we initially compared morphological RPE cell features of the human and the MG. Our results regarding RPE cell height indicate that the human RPE cell characteristics (parafoveal height of 10-15 μm vs. peripheral of about 7 μm (Feeney-Burns et al., 1990; Ishibashi et al., 2004; Weiter et al., 1986) very well match those found in the VS and periphery of the MG.

This difference between central and peripheral RPE cells in the MG was further supported by the proteomics analysis. In the RPE of the VS, we found a higher accumulation of MERTK-labelled phagosomes when compared to the peripheral retina. MERTK is a receptor present on the apical membrane of the RPE that binds indirectly to exposed phosphatidylserine of photoreceptor OSs via a soluble ligand and is necessary for the intracellular engulfment OS particles (Kevany & Palczewski, 2010; Ruggiero et al., 2012).

The upregulation of visual cycle proteins in the VS of the MG strongly suggests an increased interaction between retina and RPE in this region. This is of particular interest in conjunction with the retinal ATP-binding cassette transporter 4 (ABCA4) and the retinol dehydrogenase 8 (RDH8) enzymes, because their perturbation is responsible for increasing the levels of the toxic lipofuscin fluorophore A2E, which leads to retinal degeneration in AMD (Miller, 2013). In Rdh8^-/-^Abca4^-/-^ mice, increased levels of A2E were accompanied by RPE/photoreceptor dystrophy, for example (Maeda et al., 2008). ABCA4 is also a protein relevant for Stargardt disease (STGD1), which is the most prevalent inherited macular dystrophy and is linked to disease-causing variants in the *ABCA4* gene (Tanna et al., 2017). Taken together, we hypothesize that RPE cells in the VS, due to their increased length and volume, the increased length of photoreceptor OSs, and the increased cone photoreceptor density, encounter a higher phagocytic load and thus may be at risk for a pathological accumulation of undigested lipofuscin deposits, which we believe is a further important asset for a model of macular degeneration. Thus, the assessment of markers related to RPE and photoreceptor OS phagocytosis in the VS of (aged) MGs may be highly relevant in future studies on the pathophysiology of macular diseases.

In summary, our comparative data indicate that the MG is a promising rodent, non-primate model to investigate the pathophysiology of human central retinal and macular diseases. In terms of the grade of representation of the human macula, non-human primates actually have a macula with a fovea, which is optimal for this purpose, but their high cost, slow disease progression, difficulty for genetic manipulations and ethical concerns notably restrict their use (Pennesi et al., 2012). On the other hand, murine models have been very important to help understand key pathways of AMD disease pathology, but the lack of a macula-like region is a big limitation for this species. The MG, however, is a rodent model phylogenetically similar to mice, so that commercially available antibodies developed for mice work in their majority for the MG, but as a diurnal species its retinal organization is closer to that of larger animal models (e.g. pig, cat, dog). It shares widely the advantages of mouse models regarding low cost, short lifespan, and the availability for genetic manipulation. The latter became possible since whole-genome sequencing of the MG has been performed, and first CRISPR/Cas9-mediated mutant models have been generated (Wang et al., 2020; Zorio et al., 2019). In conclusion, the MG offers great potential as a model for the human parafovea. Still, further studies need to be performed in the future, in particular in conditions of retinal dystrophy like AMD and Stargardt disease.

## 4 Material and Methods

### Experimental animals

All animal experiments and procedures performed in this study adhered to the ARVO statement for the Use of Animals in Ophthalmic and Vision Research and were approved by the competent legal authority (Regierungspräsidium Tübingen, Germany). All efforts were made to minimize the number of animals used and their suffering. MGs were housed under an alternating 12-h light and dark cycle, with free access to food and water, and were used irrespective of gender. 5 adult MGs aged 2–3 months were used for immunohistochemical experiments and 2 MGs aged 2 years were used for proteomics analysis.

### Immunohistochemistry

The eyes from the MGs were fixed in 4% paraformaldehyde (PFA) for 45 minutes, and after that immersed in 30 % sucrose phosphate buffer (pH 7.4) overnight at 4 °C. On the next day, eyes were embedded in Tissue-Tek OCT compound (Sakura Finetek Europe, Alphen aan Den Rijn, Netherlands), slowly frozen using dry ice, and then stored at -80 °C. Retinal dorsoventral cross-sections with a 12 μm thickness were collected on Superfrost glass slides (R. Langenbrinck GmbH, SuperFrost® plus, Emmendingen, Germany) and stored at -20°C. For the immunohistochemical staining process, slides were dried at 37°C for 30 minutes and rehydrated with phosphate-buffered saline (PBS) for 10 minutes at room temperature (RT). As a next step, they were incubated for 1 hour at RT with a blocking solution that consisted of 5% chemiBLOCKER (Merck, Darmstadt, Germany) in 0.1% PBS Triton X-100. Afterwards, the sections were incubated overnight at 4 °C with the following antibodies diluted in blocking solution: FITC conjugated PNA (L7381; 1:100; Sigma-Aldrich, St. Louis, MO, USA), anti-rhodopsin (ab98887; 1:500; Abcam, Berlin, Germany) and anti-MERTK (ab95925; 1:100; Abcam). The use of a detergent in the blocking solution allowed for the visualization of intracellular MERTK-labelled phagosomes. On the next day, slides were washed 3x in a washing solution containing PBS with 2% chemiBLOCKER at RT. Then, the following secondary antibodies diluted in washing solution were incubated for 2 hours at RT: Goat anti-Rabbit, Alexa Fluor 568 (A11036; 1:300; Thermo Fischer Scientific) and Goat anti-Mouse, Alexa Fluor 647 (ab150119; 1:150; Abcam). As a last step, sections were rinsed with PBS and mounted with ROTI Mount FluorCare containing DAPI (4′,6-diamidino-2-phenylindole) (Carl Roth, Karlsruhe, Germany). DAPI was used to identify rod and RPE nuclei, and MERTK as a marker for RPE cells.

### Microscopy and image analysis

Sections used for immunohistochemistry were imaged on a Zeiss Imager Z.2 fluorescence microscope, equipped with ApoTome 2, an Axiocam 506 mono camera and an HXP-120V fluorescent lamp (Carl Zeiss Microscopy, Oberkochen, Germany). Z-stack images were captured with the ZEN 3.3 (blue edition) software (Carl Zeiss Microscopy) with a 20x magnification. The quantification of photoreceptor densities was done by averaging measurements from at least 4 sections per animal in 1200 μm^2^ sections (100 μm dorsoventral segments x 12 μm thickness) from the VS, the near and far peripheral retinas. OSs labeled with PNA were manually counted though the Z-stack to determine the density of cones. To determine the density of rods, the average diameter of outer nuclear layer (ONL) nuclei and the number of ONL rows arising from rod cells were quantified. These values were used to calculate the number of rods in 100 μm dorsoventral sections. Since the retinal sections also presented a thickness of 12 μm, the average ONL nuclei diameter was used to calculate the depth of rod cells. After multiplying these values with each other, the rod density in 1200 μm^2^ was obtained. Lastly, the photoreceptor densities calculated in 1200 μm^2^ were estimated for an area of 1 mm^2^. Dorsal and ventral sections taken approx. 300 μm from the center of the VS were considered as near peripheral retina. Dorsal sections taken approx. 2 mm from the center of the VS were considered as far periphery, as well as ventral sections with the same contralateral position. Figures were prepared using Adobe Photoshop CS5 (Adobe, San Jose, CA, USA) and the CorelDRAW X5 software (Corel corporation, Ottawa, ON, Canada). The diagram with examples of different retinal organizations was created with BioRender.com.

### Calculation of RPE cell area

For the calculation of RPE cell areas in the VS and the periphery of the MG retina, the distances between the center of RPE cell nuclei were quantified in 250 μm dorsoventral retinal segments. As some RPE cells are binucleate, we had to consider this fact in our calculations. Within the 454 RPE nuclei analysed in the VS, 64 (14%) were classified as binucleate cells, and in the peripheral retina the fraction was 73 of 703 (10%). Internuclear distances that were below half the median of the VS data (17.12 μm) or of that of the peripheral retina (20.57 μm) were considered artifacts arising from binucleate RPE cells, and distances that were above twice the medians were considered artifacts from sections where a correct intermediate nucleus was missing (4% for the VS and 5% for the peripheral retina). For binucleate RPEs, the center between the nuclei was used to calculate the distance to the next RPE cell nucleus. The quantification of distances between RPE nuclei from the above corrected data are shown in Supplementary Figure 1A.

As a next step, The RPE internuclear distances were used to calculate the resulting RPE cell areas. In detail, the calculation uses two basic formulas for equilateral triangles of side length *a* (see sketch in Supplementary Figure 1B):

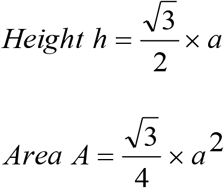

*If D is the internuclear distance, then* 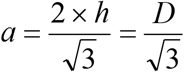, *and thus* 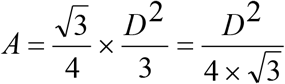

*Finally, as a hexagonconsists of* 6 *such triangles, the hexagonal area equals* 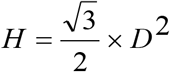

### Mass spectrometry

Eyes from MGs were enucleated, and the VS was isolated with a sharp scalpel under a dissection microscope. The VS appears as a light horizontal band visible in the retina. Accordingly, a dorsal region lying adjacently from the VS was isolated with the same size as the former one. These regions were sent for mass spectrometry as the intact RPE + retina, or as the isolated RPE. The analysis was performed on an Ultimate3000 RSLC system coupled to an Orbitrap Tribrid Fusion mass spectrometer (Thermo Fisher Scientific). Tryptic peptides were loaded onto a µPAC Trapping Column with a pillar diameter of 5 µm, inter-pillar distance of 2.5 µm, pillar length/bed depth of 18 µm, external porosity of 9%, bed channel width of 2 mm and length of 10 mm; pillars are superficially porous with a porous shell thickness of 300 nm and pore sizes in the order of 100 to 200 Å at a flow rate of 10 µl per min in 0.1% trifluoroacetic acid in HPLC-grade water. Peptides were eluted and separated on the PharmaFluidics µPAC nano-LC column: 50 cm µPAC C18 with a pillar diameter of 5 µm, inter-pillar distance of 2.5 µm, pillar length/bed depth of 18 µm, external porosity of 59%, bed channel width of 315 µm and bed length of 50 cm; pillars are superficially porous with a porous shell thickness of 300 nm and pore sizes in the order of 100 to 200 Å by a linear gradient from 2% to 30 % of buffer B (80% acetonitrile and 0.08% formic acid in HPLC-grade water) in buffer A (2% acetonitrile and 0.1% formic acid in HPLC-grade water) at a flow rate of 300 nl per min. The remaining peptides were eluted by a short gradient from 30% to 95% buffer B; the total gradient run was 120 min.

Spectra were acquired in DIA (Data Independent Acquisition) mode using 50 variable-width windows over the mass range 350-1500 m/z. The Orbitrap was used for MS1 and MS2 detection, with an AGC target for MS1 set to 20×10^4^ and a maximum injection time to 100 ms. MS2 scan range was set between 200 and 2000 m/z, with a minimum of 6 points across the peak. Orbitrap resolution for MS2 was set to 30K, isolation window set to 1.6, AGC target to 50×10^4^ and maximum injection time to 54 ms. MS1 and MS2 data were acquired in centroid mode. To reduce the possibility of carry over and cross contamination between the samples, one TRAP and two BSA washes were used between samples groups.

### Statistical analysis

A two-tailed paired student’s t-test and a repeated-measures one-way ANOVA followed by Bonferroni post hoc test was used to assess statistical differences between quantitative data of two and three groups, respectively. All analyses were based on five individual animals. Values of p < 0.05 were considered to be statistically significant and labeled with an asterisk (*) in the graphs. To indicate a higher degree of statistical significance, values of p < 0.01 were marked with two (**) asterisks and p < 0.001 with three (***) asterisks. Graphical results are represented as mean ± standard deviation for photoreceptor density and normalized RPE length metrics. In boxplots, used to display ratios, distance between RPE cell nuclei and RPE cell areas, the limits represent the 25%-75% quantile range, the central line the median, and the whiskers designate 5% and 95% quantiles. The statistical analysis was performed with GraphPad Prism 10.1 for Windows (GraphPad Software, La Jolla, CA, USA).

MS RAW data were analysed using DIA-NN 1.8.2 beta 27 (PMID: 31768060) in library-free mode against the NCBI Gerbil database (release December 2023, 21261 proteins). First, a precursor ion library was generated using FASTA digest for library-free search in combination with deep learning-based spectra prediction. An experimental library generated from the DIA-NN search was used for cross-run normalisation and Mass accuracy correction. Only high-accuracy spectra with a minimum precursor FDR of 0.01, and only tryptic peptides (2 missed Tryptic cleavages) were used for protein quantification. The match between runs option was activated and no shared spectra were used for protein identification.

Data where further analysed using the Perseus platform (Tyanova et al., 2016). First, data were log2-transformed to facilitate the identification of proportional changes in metabolite abundance between the different groups. This transformation helps normalize the data and stabilize the variance, making it more suitable for subsequent statistical analysis. To determine significant differences in proteins abundance between groups, a two-sample t-test was employed. This test was complemented by a permutation-based false discovery rate (FDR) control method, set at a threshold of 0.05, to account for multiple comparisons and reduce the likelihood of type I errors. Specifically, 250 random permutations of the data were performed to generate the null distribution and calculate the FDR-adjusted p-values. This approach ensures robust identification of statistically significant changes in proteins abundance.

## Ethics

The animal study protocol was approved by the competent animal ethics committee of the regional authorities (Regierungspräsidium Tübingen).

## Author contributions

Alexander Günter: Conceptualization, data curation, formal analysis, investigation, methodology, visualization, writing – original draft preparation. Mohamed Ali Jarboui: Validation, formal analysis, investigation, data curation, Writing – review & editing. Regine Mühlfriedel: Funding acquisition, project administration, resources, writing – review & editing. Mathias Seeliger: Conceptualization, methodology, funding acquisition, project administration, resources, supervision, writing – review & editing.

## Acknowledgements

We thank Gudrun Utz for the excellent technical assistance. Figure 1 was created with BioRender (https://biorender.com accessed on 26 August 2024).

