## Supplementary Figures for "A remarkable degree of conformance between the visual streak of the Mongolian gerbil and the human central retina"

**This file includes supplementary Figures S1 and S2.**

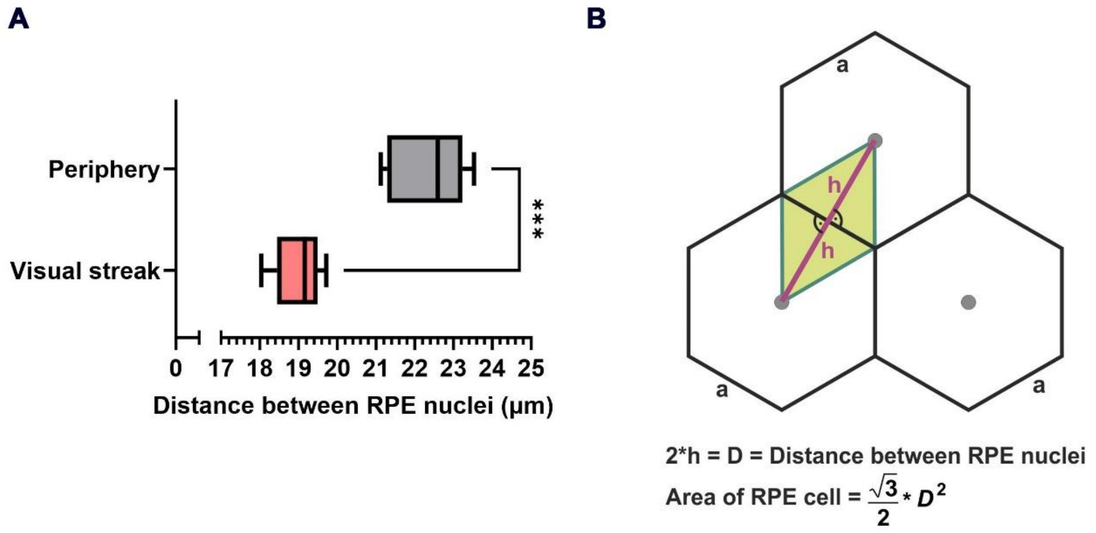


**Figure S1. Parameters for the analysis of RPE cell areas. (A)** Quantitative evaluation of the distances between the RPE nuclei (box-and-whisker-plot) in the VS (n = 5) compared to the peripheral retina. Boxes: 25%-75% quantile range, whiskers: 5% and 95% quantiles, central line: median. **(B)** Mathematical model based on the simplifying assumption of a hexagonal RPE monolayer used to calculate RPE cell area based on the distances between nuclei. D = distance between nuclei; h = height; a = side.

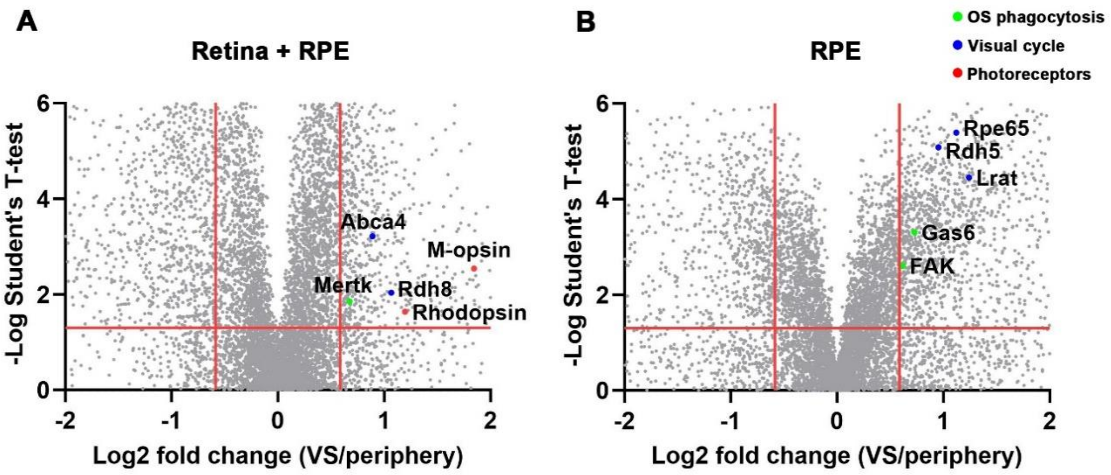


**Figure S2. Proteomic mass spectrometry-based differential expression analysis of MGs in the VS compared to the peripheral retina.** Volcano plots showing significant differential abundant proteins in **(A)** the interface of the retina + RPE and **(B)** in the isolated RPE by quantitative proteomic analysis. The p-value was plotted on the y-axis as $-\log_{10}$ and the relative abundance ratio of VS/periphery on the x-axis as a $\log_2$ fold change. Red vertical lines indicate the cutoff for a significant enrichment- or depletion of protein abundance of 1.5-fold. The red horizontal line indicates a p-value of 0.05. VS = Visual streak.
